# Describing macroecological patterns in microbes: Approaches for comparative analyses of operational taxonomic unit read number distribution with a case study of global oceanic bacteria

**DOI:** 10.1101/778829

**Authors:** Ryosuke Nakadai, Yusuke Okazaki, Shunsuke Matsuoka

## Abstract

Describing the variation in commonness and rarity in a community is a fundamental method of evaluating biodiversity. Such patterns have been studied in the context of species abundance distributions (SADs) among macroscopic organisms in numerous communities. Recently, models for analyzing variation in local SAD shapes along environmental gradients have been constructed. The recent development of high-throughput sequencing enables evaluation of commonness and rarity in local communities of microbes using operational taxonomic unit (OTU) read number distributions (ORDs), which are conceptually similar to SADs. However, few studies have explored the variation in local microbial ORD shapes along environmental gradients. Therefore, the similarities and differences between SADs and ORDs are unclear, clouding any universal rules of global biodiversity patterns. We investigated the similarities and differences in ORD shapes vs. SADs, and how well environmental variables explain the variation in ORDs along latitudinal and depth gradients. Herein, we integrate ORD_S_ into recent comparative analysis methods for SAD shape using datasets generated on the Tara Oceans expedition. About 56% of the variance in skewness of ORDs among global oceanic bacterial communities was explained with this method. Moreover, we confirmed that the parameter combination constraints of Weibull distributions were shared by ORDs of bacterial communities and SADs of tree communities, suggesting common long-term limitation processes such as adaptation and community persistence acting on current abundance variation. On the other hand, skewness was significantly greater for bacterial communities than tree communities, and many ecological predictions did not apply to bacterial communities, suggesting differences in the community assembly rules for microbes and macroscopic organisms. Approaches based on ORDs provide opportunities to quantify macroecological patterns of microbes under the same framework as macroscopic organisms.

## Introduction

Species diversity patterns, which are characterized by numbers of species and individuals, provide great opportunities for understanding the ecological and evolutionary processes that drive global biodiversity (Rabinowitz, 1981; Ricklefs, 2000; Hubbell, 2001, 2013; Loza *et al*., 2017). In general, certain species are dominant while others are rare locally and globally. The processes determining locally common and rare species have been studied in the context of the species abundance distribution (SAD), which is one of the most fundamental methods of describing local diversity (Motomura, 1932; MacArthur, 1960; McGill *et al*., 2007; Doi and Mori, 2013; Ulrich *et al*., 2018b). SAD studies focus on the rarity and commonness of species in a local community, and attempt to reconstruct the background processes that led to those patterns. To date, numerous SAD models have been proposed (reviewed in McGill *et al*., 2007). MacArthur (1957, 1960) developed the broken stick model based on the hypothesized niche portioning process, and Hubbell *et al*. (2001) provided a mechanistic interpretation of observed abundance distributions with well-defined ecological parameters such as dispersal, speciation rate, local abundance, and meta-community size, under the premise of ecological drift. Using these models, researchers estimated background processes from SAD patterns. Previous SAD investigations have been conducted mainly in plants and animals (e.g., Ulrich *et al*., 2010; Baldridge *et al*., 2016), because large datasets and specific criteria for species delimitation are necessary for SAD. Therefore, the SAD approach has been applied less to studies of microorganisms than those of macroscopic species.

The recent development of molecular techniques, in particular high-throughput sequencing, has made it dramatically easier to capture biodiversity patterns in microbes as well as larger organisms (Lynch and Neufeld, 2015; Schloss *et al*., 2016; Shade *et al*., 2018). High-throughput sequencing technology has revealed that the microbial ecosystem is inhabited by a large number of rare microbial lineages (Fuhrman, 2009), collectively referred to as the “rare biosphere” (Sogin *et al*., 2006; Galand *et al*., 2009; Ser-Giacomi *et al*., 2018; reviewed in Lynch and Neufeld, 2015). The existence of rare lineages strongly impacts community structure and diversity patterns, although their ecological roles and the processes structuring the rare biosphere remain poorly understood (Ser-Giacomi *et al*., 2018). The relationships between operational taxonomic units (OTUs; reviewed in Blaxter *et al*., 2005) and read number have been studied in microbes based on concepts similar to SAD (e.g., Livermore and Jones, 2015). Hereafter, we consider OTU read number distributions (ORDs), corresponding conceptually to the rank abundance curve, to be distinct from SADs. OTUs are generally defined as groups based on sequence similarity in maker genes (e.g., SSU rRNA in bacteria and ITS in fungi) (Bálint *et al*., 2016). Numerous previous studies have attempted to identify a general best-fit model for ORD in a particular habitat by comparing various traditional SAD models (e.g., log-series, lognormal, and power-law distributions) (Shade *et al*., 2012, Sherrill-Mix *et al*., 2016, Shoemaker *et al*., 2017, Louca *et al*., 2019). Only a few studies have focused on the continuous variation in ORD shape within microbial communities along geographical and environmental gradients (Stegen *et al*., 2016), which has been a major recent trend in SAD studies of macroecological patterns and their assembly processes.

In recent years, methods for analyzing macroecological patterns and community assembly by comparing SAD shapes have been developed to overcome the limitations of conventional SAD modeling. Specifically, researchers have recently recognized that multiple processes can generate similar SAD shapes, resulting in the fit of a given SAD model not providing clear evidence to support a particular theory (Mathews *et al*., 2017). An alternative approach to comparing changes in parameter value(s) for a given SAD model or the fits of different SAD models to data from a single site is the collection of abundance data from a variety of sites followed by construction of models to analyze how SAD properties vary with predictor variables (i.e., comparative analyses of SAD shape; White *et al*., 2012, Ulrich *et al*., 2016a, b, Fattorini *et al*., 2016; Borda-de-Água *et al*., 2017, Guerin *et al*., 2017, Mathews *et al*., 2019; reviewed in Mathews *et al*., 2017). Many methods have been developed to evaluate SAD shape. For example, ρ_norm_ has been used to evaluate the relative fits of log-series and lognormal distributions (Ulrich *et al*., 2016 a, b). However, the fits of both log-series and lognormal distributions to community data often weaken with increasing species richness (Ulrich *et al*., 2016a, b), which makes the interpretation of these fits inconsistent along species richness gradients in community data. The gambin distribution (Ugland *et al*., 2007) is often used to evaluate SAD shape (Mathews *et al*., 2014, 2017, 2019), but creation of the gambin distribution requires binning the abundance data into octaves prior to fitting, losing species-level abundance information. In addition, the current criterion used to fit the model is the chi-square test, which generally has a strong dependence on sample size, which makes it difficult to apply the gambin distribution go high-throughput sequencing data with large numbers of both OTUs and reads.

The comparative analysis approaches used for SAD shape provide two new insights. First, estimating the constraints of community assembly involves analyzing macroecological patterns. Ulrich *et al*. (2018b) proposed the concept of ‘forbidden communities’ in SAD shapes, placing limitations on the possible combinations of λ (scale parameter) and η (shape parameter), based on the analytical results of global tree community data. The parameter λ can be interpreted as a measure of SAD shape-specific evenness. The shape parameter η is associated with an excess of either highly abundant species (low η) or rare species (high η). Specifically, Ulrich *et al*. (2018b) argued that the combination of high η and low λ generates SADs very similar to the well-known broken stick distribution (MacArthur 1957), but those distributions rarely occur in nature. Ulrich *et al*. (2018b) showed that the shape and scale parameters of this distribution have precise ecological interpretations, with the first acting as a measure of the excess of either rare or common species and the second quantifying the proportion of persistent species in the local community. In addition, Ulrich *et al*. (2018b) showed that the scale parameter is linearly correlated with failure time mathematically, and that it indicates the relative proportion of adapted species in a community that may be persistent. Thus, these forbidden communities may indicate limited long-term processes such as adaptation and the persistence of the community characterized by the current abundance variation.

Recently, biogeographic and macroecological studies on geographical and climatic variation in SADs have employed large datasets, mainly for plants (White *et al*., 2012; Ulrich *et al*., 2016a, b; Guerin *et al*., 2017). Indeed, large-scale SAD studies have recently gained attention due to the accessibility of databases. A large amount of species abundance data collected across a broad range of environments is available for large-scale comparative SAD research (Fattorini *et al*., 2016; Borda-de-Água *et al*., 2017). For example, Ulrich *et al*. (2016a) generated a spectrum of lognormal to log-series type distributions in relation to deterministic niche differentiation vs. stochastic dispersal-driven processes along environmental climatic and productivity gradients. Thus, Ulrich *et al*. (2016a) showed latitudinal patterns of SAD shape, which is an excess of log-series SADs at lower latitudes. This suggested that the importance of dispersal-driven processes for local tree community assembly increases toward lower latitudes, which are higher productivity sites, and *vice versa*. In another example, Guerin *et al*. (2017) reported that geographical and climatic gradients affect SAD shapes and discussed future changes in their shapes expected as a consequence of climate change, with changes in diversity and ecosystem function.

Here, we integrate recent approaches used for comparative analyses of SAD shape into ORDs. As a case study, we used community datasets generated form the Tara Oceans expedition (Sunagawa *et al*., 2015). Tara Oceans is a project that profiled planktonic and microbial communities in the global ocean using high-throughput sequencing. We investigated potential factors affecting variation in ORDs from local bacterial communities along a geographic gradient, and assessed the similarities and differences between the geographical patterns of microbes and macroscopic organisms, comparing the ORDs of bacterial communities with those of macroscopic communities. In particular, we tested three hypotheses for determining skewness in microbial communities based on existing studies in macroscopic organisms. First, climatic and geographic factors, which are directly associated with local productivity, drive skewness (productivity hypothesis), such that higher productivity facilitates lower skewness (Whittaker, 1975; Hubbell, 1979; Ulrich *et al*., 2016a). Second, local concentrations of nutrients in water negatively affect skewness (nutrient hypothesis); thus, a more eutrophic environment exhibits lower skewness. Finally, characteristics of microbial lifecycle strategies (i.e., longevity and persistence) negatively influence skewness (r-K strategy hypothesis), such that shorter generation times lead to greater skewness. Detailed explanations of those hypotheses and predictions are provided in Table 1. In addition, we studied the similarities and differences in ORD and SAD shapes, with a particular focus on skewness and parameter combinations of the Weibull distribution, and the extent to which environmental variables explain the variation in ORDs along latitudinal and depth gradients. This approach could support further acceleration of microbial studies by revealing the drivers of variation in ORD shape and making comparisons between microbes and macroscopic species.

**Table 1.** Major hypotheses and predictions associated with skewness tested in this study

| Hypothesis | Prediction | Related variables | References |
| --- | --- | --- | --- |
| (i) Low productivity conditions facilitate skewness (productivity hypothesis). | Skewness decreases toward lower-latitude, deeper, and cooler conditions. | Latitude, Water depth, Temperature | Whittaker (1975), Hubbell (1979), Ulrich *et al.* (2016a) |
| (ii) Nutrient limitation facilitates skewness (nutrient hypothesis). | Skewness increases toward lower-nutrient conditions. | NO<sub>2</sub>, PO<sub>4</sub> | Ulrich *et al.* (2016a) |
| (iii) Dominance of r strategists facilitates skewness (r-K strategy hypothesis). | Skewness decreases with increasing long-lived OTUs. | Minimum potential generation time | Magurran & Henderson (2003), Ulrich & Ollik (2004), Ulrich *et al.* (2016a) |

## Materials and Methods

### Empirical data

We used 139 oceanic bacterial community datasets generated from the Tara Oceans expedition (Sunagawa *et al*., 2015). Sunagawa *et al*. (2015) extracted merged metagenomic Illumina reads (miTAGs) containing signatures of the 16S rRNA gene (Logares *et al*., 2013), which were mapped to the SILVA SSU rRNA gene sequence database (Quast *et al*., 2013) and clustered into OTUs at the 97% similarity level. The range of total reads per sample was 34,081–184,190, with an average of 90,103.06 ± 28,240.38. The OTU count table was summarized at multiple taxonomic levels and can be downloaded from http://ocean-microbiome.embl.de/companion.html (Sunagawa *et al*., 2015). We extracted OTUs representing the domain Bacteria for analyses.

We calculated the skewness *γ* of log-transformed relative read numbers to assess the degree of lower curvature (Ulrich *et al*., 2016a), which was compared to a symmetrical lognormal distribution. Negative values of *γ* indicate an excess of rare OTUs, while positive values represent an excess of abundant OTUs compared to a lognormal distribution that is symmetrical around the mean. Thus, the symmetrical lognormal distribution is not skewed. Asymmetrical lognormal SADs nearly always indicate an excess of rare OTUs, and consequently have negative skewness (McGill, 2003). The log-series model shows an excess of relatively abundant OTUs (associated with positive skewness). We used the parametric function SE (*γ*) = (6/n)^1/2^ (Tabachnick and Fidell, 1996) to test for significant skewness in SADs (Ulrich *et al*., 2016a). In this bacterial community dataset, skewness is positively correlated with the number of rare OTUs (rare biosphere: <0.1 or <0.01, criteria based on Galand *et al*., 2009, Pedrós-Alió *et al*., 2012) (Fig. S1). The relationships among ρ_norm_, the alpha parameter of the gambin distribution, and skewness are shown in Fig. S3.

### Comparison between ORDs of microbes and SADs of forest plots

To reveal differences between the ORD shapes of microbes and SAD shapes of macroscopic organisms, we first compared the skewness values of bacterial communities with those of tree communities published in Appendix 1 of Ulrich *et al*. (2016a). In addition, we compared the two-parameter Weibull distribution, which was recently suggested by Ulrich *et al*. (2018b) as a flexible descriptive model for SAD shapes. The empirical Weibull distribution (Weibull, 1951) is an extension of the exponential family of distributions (Rinne, 2008) and has been widely used in survival analyses (Lawless, 2003). Stauffer (1979) proposed use of the Weibull distribution as a model to explain observed species abundance distributions (Ulrich et al. 2018b). Ulrich *et al*. (2018b) suggested that the shape and scale parameters of the Weibull distribution have precise ecological interpretations, with the first being a measure of the excess of either rare or common species, and the second quantifying the proportion of persistent species in the focal community. To identify any similar limitations of parameter combinations in bacterial communities, we fitted two-parameter Weibull distributions to datasets of oceanic bacterial communities and compared the ORD shapes for microbes with the empirical results from 534 tree communities worldwide, presented in Ulrich *et al*. (2018a). If the ORDs of bacterial communities show similar constraints, community structuring processes are likely shared between the bacterial and tree communities, although the specific mechanisms are unknown. We used the reduced major axis fitting method, following Ulrich *et al*. (2018b). Values of *fit* < 0.05 indicate an excellent fit, while *fit* > 0.3 is poor. We used the updated stand-alone application RAD 2.0 (Ulrich *et al*., 2010) for fitting of the two-parameter Weibull distributions.

### Explanatory variables for skewness in ORDs

In the multi-regression model, we used measurements provided in the original dataset from Sunagawa *et al*. (2015). We set latitude, squared latitude, water depth (m), nitrite concentration (µmol L^-1^), phosphate concentration (µmol L^-1^), and minimum potential generation time (h) as explanatory variables for the variation in skewness. The average minimum potential generation times of microbial communities, which were determined from codon usage biases in metagenome sequences (for detailed methods, see Vieira-Silva and Rocha 2010), were used as an index of r-K strategy at the community level.

The data output from high-throughput sequencing generally contains different sequencing depths (i.e., uneven sampling effort) for each sample. Sampling effort influences observed SAD shape (Preston, 1948). Therefore, to account for the effects of biases in the sampled data, we added the OTU richness and total read number of each sample as covariates in the regression models. To avoid nonlinear effects, we used ln-transformed data for water depth, species richness, and total abundance.

The level of collinearity between these explanatory variables was determined by calculating the variance inflation factor (VIF). All variables were standardized to zero mean and unit variance prior to parameter estimation. We selected explanatory variables according to a threshold VIF value, where VIF > 10 indicates that the model has a collinearity problem (Quinn and Keough, 2002, Neves *et al*., 2015). VIF ranged from to 8.26, suggesting a lack of multicollinearity in this multi-regression model. However, the effects of temperature on skewness showed the opposite trends compared to the results of simple correlation (Fig. S4). We excluded temperature from the final analyses, causing the VIFs to range from 1.07 to 3.06. We use the ‘car’ package of R (Fox *et al*., 2007) to calculate VIFs and the ‘rsq’ package (Zhang, 2018) to calculate partial r-square coefficients.

## Results

The mean of skewness was 0.81±0.17 among 139 samples. All communities showed significant positively skewed trends compared to the symmetric lognormal model (Tabachnick and Fidell, 1996; Ulrich *et al*., 2016a), indicating that all ORD shapes are significantly more similar to log-series distributions than lognormal distributions. Based on the results of t-test, bacterial communities are significantly more skewed than tree communities (Welch’s t-test; p<0.0001), and thus more similar to the log-series distribution in shape.

Weibull distribution fits to 137 of the 139 communities (98.6%) were moderate (fit < 0.3), while only two (1.4%) were comparatively poor (fit>0.3). The parameter combinations in bacterial communities did not exceed the parameter space for tree communities (Fig. 1), suggesting constraints on ORD shape (Ulrich *et al*., 2018a).

**Figure 1.**
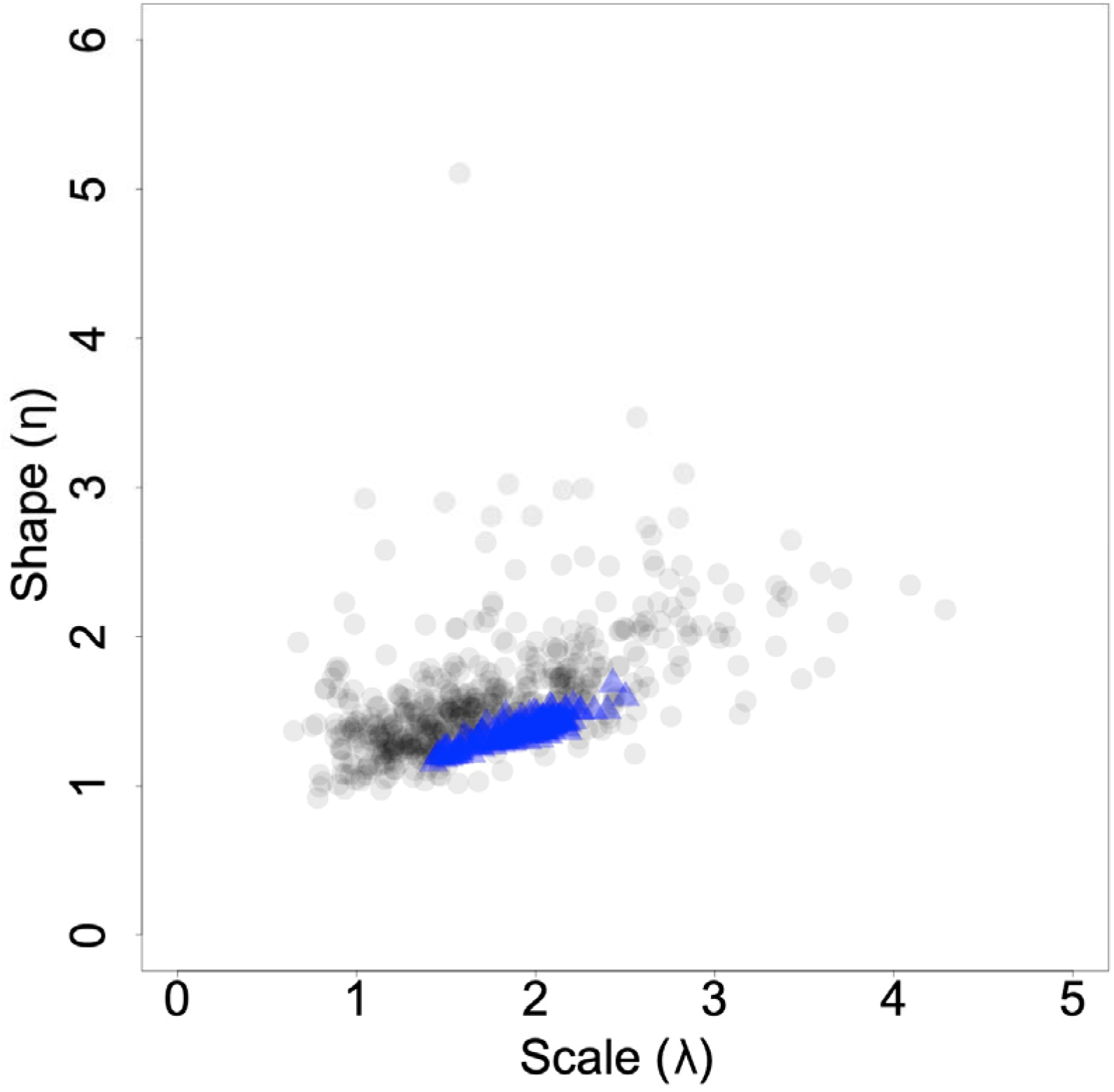
Relationship between shape (η) and scale (λ) parameters. Blue circles indicate the parameters of 139 bacterial communities analyzed in this study. Gray circles indicate the parameters of 534 empirical global tree communities published in Ulrich *et al*. (2018a) and https://figshare.com/articles/Weibull_fits/5975098. The parameters of bacterial communities did not exceed the parameter space of tree communities.

In total, the model explained about 56% of the variance in skewness in 131 ORDs (Table 2). Eight communities lacked some data and were excluded from the analyses of ORD shapes. Skewness increased with increasing minimum potential generation time, and was not associated with productivity, nutrients or r-K strategy, as shown in Table 1.

**Table 2.** Results of the multi-regression model for skewness.

| | $\beta$ | $r^2_{par}$ | $P$ | |
| --- | --- | --- | --- | --- |
| Latitude | -0.0425 | 0.0034 | 0.5190 |  |
| Latitude<sup>2</sup> | 0.0817 | 0.0150 | 0.1740 |  |
| ln(Water depth) | 0.0304 | 0.0007 | 0.7660 |  |
| NO<sub>2</sub> | 0.0530 | 0.0064 | 0.3770 |  |
| PO<sub>4</sub> | 0.1397 | 0.0120 | 0.1180 |  |
| Minimum potential generation time | 0.3884 | 0.1634 | <0.0001 | *** |
| ln(Total read number) | 0.0190 | 0.0006 | 0.7830 |  |
| ln(OTUs richness) | 0.3863 | 0.1335 | <0.0001 | *** |
| Whole model | | $adj. R^2$ | $P$ | |
|  |  | 0.5591 | <0.0001 | *** |

Significance: \*<0.05, \*\*<0.01, \*\*\*<0.001

## Discussion

We fit the two-parameter Weibull distribution and clearly showed that the parameters of the bacterial communities were not outside of the limited space described in Ulrich *et al*. (2018b), suggesting some limitation on community assembly processes that is shared between SADs and ORDs. We could explain about 56% of skewness variation in ORDs, revealing potential factors affecting skewness. Therefore, we conclude that comparative analyses of ORDs shape are also useful for identifying potential affecting factors and hypothesized background processes, and that ORD shapes have similar constraints to those described in Ulrich *et al*. (2018b). These findings emphasize the applicability of comparative analyses of ORD shape in microbial communities. However, the number of case studies using this method remains small, so further empirical studies are needed to fully elucidate ORD patterns and their drivers.

### Similarities and differences between SADs and ORDs

Comparison of SADs and ORDs has been suggested as a useful approach to identify general rules across microbial and macroscopic communities (Shade *et al*., 2018). In the present study, we compared SADs of tree communities with ORDs of microbial communities using the two fitted parameters (shape parameter [η] and scale parameter [λ]) of the Weibull distribution. We confirmed similar constraints of parameter combinations in ORDs and SADs of tree communities of η < 3 and λ < 6, as originally proposed in Ulrich *et al*. (2018b). These constraints on community structure may indicate limitation due to long-term processes such as adaptation and persistence of the community on variation in abundance (Ulrich *et al*. 2018b). In the present study, we defined persistent species/OTUs as those that were abundant and biologically associated with suitable habitats (Magurran and Henderson 2013). In contrast, we defined transient species/OTUs as those with low abundance and different habitat requirements (Magurran and Henderson 2013). Furthermore, we defined community persistence and transience as the relative dominance proportions of persistent/transient species within each community (Ulrich et al. 2018b). Locey and Lennon (2016) found similar scaling of commonness and rarity across microbes and macroscopic plants and animals, and proposed a universal dominance scaling law that holds true over 30 orders of magnitude. We suggest that the concept of forbidden communities presented by Ulrich *et al*. (2018b) is related to the scaling law (e.g., community persistence is regulated by metabolic rate). Further comparisons of SADs and ORDs may reveal whether similar constraints exist in other taxonomic groups and habitats.

We emphasize that ORDs should have similar dataset properties as SADs. We also have a negative view regarding the common practice of referring to ORDs as SADs and their inclusion in SAD studies. In our analyses, the skewness of bacterial communities was significantly greater than that of tree communities, indicating a larger proportion of rare OTUs among bacteria than among trees. Previous studies in macroscopic organisms have reported that persistent species (i.e., stable communities driven mainly by habitat filtering) exhibit a lognormal SAD (i.e., smaller skewness), while transient species (i.e., dispersal-driven communities that vary over time) are best modeled with the log-series (i.e., high skewness) distribution (Magurran and Henderson, 2003; Ulrich and Ollik, 2004; Ulrich *et al*., 2010, 2016a). Therefore, our results suggest that bacterial communities could include larger proportions of transient OTUs if we interpret the results based on findings from macroscopic organisms. Because time-series data were unavailable in the present study, this hypothesis should be tested in future studies using temporal datasets.

There are two major factors that should be considered with respect to the inclusion of ORDs in SAD studies. The first issue is associated with delimitation of species and individuals in microbes (reviewed in Hason *et al*., 2012). Empirically, 97% similarity in (partial) 16S rRNA gene sequences has commonly been used to characterize prokaryotic species-level phylogenetic diversity. However this resolution can lead to underestimation of genomic (i.e., species) diversity (Rodriguez-R *et al*., 2018). In tree communities, Hubbell (2013) found differences in SAD shape among taxonomic resolutions (i.e., species, genus, and family) in tropical forest data; specifically, the tail length (rare species) decreased with lower taxonomic resolution. Moreover, researchers generally treat the read number of OTUs as an abundance metric. However, the results of 16S rRNA gene sequencing reveal relative, rather than absolute, abundances of individual lineages. In addition, these results may be biased due to differences in PCR primer specificity and gene copy number among lineages. Thus, care should be taken when comparing those patterns with actual abundance data.

The second issue is associated with the community characteristics. In SAD studies, researchers generally focus on a horizontal community composed of a single trophic level, such as “tree community” and “herbivore community” (but see Mathews *et al*., 2019). However, the sequencing data based on a marker gene contains DNA of the entire assemblage in a sample, which makes it impossible to categorize the trophic levels of all microbes and ontogenetic stages. The presence of a persistent microbial seed bank (i.e., *in situ* populations of long-lived rare OTUs) might affect community-level patterns (Gibbsons *et al*., 2013), making it difficult to interpret the relationship between the proportion of rare OTUs and minimum potential generation time. In other words, microbial community datasets include different life stages as one category, analogous to grouping “seed in the soil,” “mature tree,” and “dead tree” in a dataset of trees. Above, we noted the possibility that bacterial communities include larger proportions of transient OTUs when interpreted based on data for macroscopic organisms. If we consider that long-lived rare OTUs are present *in situ*, the interpretation of a larger proportion of transient OTUs in microbes is directly contradicted. Therefore, it may be difficult to apply ecological rules that were developed in horizontal communities of macroscopic species (Vellend *et al*., 2010, Vellend 2016) to DNA-based microbial datasets, at least those related to the community properties of persistence and transience.

### Extent to which environmental variables explain variation in ORD shapes in oceanic bacteria worldwide

The multi-regression model explained about 56% of skewness variation in ORDs. Using data from oceanic bacterial communities, we confirmed that the patterns predicted from hypotheses based on macroscopic studies were not supported, as shown in Table 1. Furthermore, the skewness was significantly and positively affected by the minimum potential generation time within bacterial communities. This result is in opposition to the prediction based on research in macroscopic organisms (Table 1, 2). To explain the opposing trends in bacterial communities compared to predictions for macroscopic species, we could interpret this result as showing that competitive exclusion is less likely occur in bacteria than in macroscopic communities. In macroscopic organisms, species are adapted to the environment as a community, which makes the relationship between the environment and community structure clear. In animals and plants, when suitable species are present in an environment, unsuitable species are excluded from the community, and the relationship between the environment and community structure is simple. On the other hand, some bacteria under unsuitable conditions may continue existing (e.g., enter dormancy), thus appearing in high-throughput sequencing data. Campbell *et al*. (2011) compared DNA-based patterns with those based on RNA (i.e., transcriptionally active OTUs), and suggested that about 12% of amplicon sequences of oceanic bacteria are always inactive. Falkowski and Oliver (2007) previously argued that the distributional patterns of marine phytoplankton could be explained using a mathematical model based on the resource competition theory. Bacteria sharing resources, such as organic matter derived from phytoplankton, would be affected by competitive exclusion (Falkowski and Oliver 2007); therefore, testing the applicability of our interpretation in other microbial communities remains an important issue for future studies.

### Conclusions and perspectives

Comparative analyses of skewness, ρ_norm_, and the Weibull and gambin parameters, which are major approaches used recently in SADs, provide researchers a basis for discussing similarities and differences between microbial and macroscopic life. In the near future, comparative approaches between microbes and macroscopic organisms, including environmental DNA (eDNA) metabarcoding studies (e.g., Doi *et al*., 2019), may reveal universal rules determining global biodiversity patterns. Further theoretical frameworks focused specifically on microbes and multi-trophic data are needed, with macroscopic ecology studies such as Hubbell *et al*. (2001) as a starting point for discussion (Rosindell *et al*., 2012). We encourage construction of microbe-specific ecological rules, such as rules explicitly considering the metabolic versatility of microbes in community assembly processes, as the properties necessary for inclusion in datasets differs from those for macroscopic organisms. At the same time, further comparative analyses may reveal the detailed drivers and facilitate a better understanding of the similarities and differences between the quantitative patterns of macroscopic and microbial communities.

## Acknowledgments

We thank Dr. Doi Hideyuki for providing fruitful comments on an early version of our manuscript, and the handling editor and one anonymous reviewer for their comments which improved our manuscript. Financial support was provided by the Japan Society for the Promotion of Science (No. 18J00093 to RN).

## Data Accessibility Statement

All environmental factors, calculated data, and the OTU table used for analyses are presented in Tables S1 and S2, and the original data were published in Sunagawa *et al*. (2015) and http://ocean-microbiome.embl.de/companion.html. The Weibull parameters and skewness of global tree communities published in Ulrich *et al*. (2018a) are available from https://figshare.com/articles/Weibull_fits/5975098 and Appendix 1 of Ulrich *et al*. (2016a), respectively.

---

Figure S1 Positive correlations between skewness and rare OTU richness ((a) *adj*. *R*^*2*^ = 0.3951, *p* <0.0001, (b) *adj*. *R*^*2*^ = 0.3966, *p* <0.0001).

Figure S2 Relationship between lognormal and log-series fits ((a) *adj*. *R*^*2*^ = –0.0045, *p* = 0.5387, (b) *adj*. *R*^*2*^ = 0.1243, *p* < 0.0001, (c) *adj*. *R*^*2*^ = 0.0840, *p* = 0.0003). We followed the methods of Ulrich *et al*. (2016a) for calculating both log-series and lognormal fits.

Figure S3 Comparison among ρ_norm_, gambin alpha, and skewness. We calculated two indices used for comparative analyses of SAD shapes, ρ_norm_ and gambin alpha. ((a) *adj*. *R*^*2*^ = 0.3173, *p* <0.0001, (b) *adj*. *R*^*2*^ = 0.4440, *p* <0.0001, (c) *adj*. *R*^*2*^ = 0.7773, *p* <0.0001). We followed the methods of Ulrich *et al*. (2016a) and Mathews *et al*. (2014) to calculate ρ_norm_ and gambin alpha, respectively.

Figure S4 Results of simple correlation testing. Significance: *<0.05, **<0.01, ***<0.001, ^■^<0.1

Table S1 Extracted bacterial OTU table published in Sunagawa et al. 2015.

Table S2 Environmental variables and calculated indices used in this study. Supplementary text 1 Multimodality of the gambin distribution.

## References

Baldridge, E., Harris, D. J., Xiao, X., & White, E. P. (2016). An extensive comparison of species-abundance distribution models. PeerJ, 4, e2823.

Bálint, M., Bahram, M., Eren, A. M., Faust, K., Fuhrman, J. A., Lindahl, B., … & Tedersoo, L. (2016). Millions of reads, thousands of taxa: microbial community structure and associations analyzed via marker genes. FEMS microbiology reviews, 40(5), 686–700.

Blaxter, M., Mann, J., Chapman, T., Thomas, F., Whitton, C., Floyd, R., & Abebe, E. (2005). Defining operational taxonomic units using DNA barcode data. Philosophical Transactions of the Royal Society B: Biological Sciences, 360(1462), 1935–1943.

Borda-de-Água, L., Whittaker, R. J., Cardoso, P., Rigal, F., Santos, A. M., Amorim, I. R., … & Borges, P. A. (2017). Dispersal ability determines the scaling properties of species abundance distributions: a case study using arthropods from the Azores. Scientific reports, 7(1), 3899.

Campbell, B. J., Yu, L., Heidelberg, J. F., & Kirchman, D. L. (2011). Activity of abundant and rare bacteria in a coastal ocean. Proceedings of the National Academy of Sciences, 108(31), 12776–12781.

Doi, H., & Mori, T. (2013). The discovery of species–abundance distribution in an ecological community. Oikos, 122(2), 179–182.

Doi, H., Inui, R., Matsuoka, S., Akamatsu, Y., Goto, M., & Kono, T. (2019). Evaluation of biodiversity metrics through environmental DNA metabarcoding outperforms visual and capturing surveys. bioRxiv, 617670.

Fattorini, S., Rigal, F., Cardoso, P., & Borges, P. A. (2016). Using species abundance distribution models and diversity indices for biogeographical analyses. Acta oecologica, 70, 21–28.

Falkowski, P. G., & Oliver, M. J. (2007). Mix and match: how climate selects phytoplankton. Nature reviews microbiology, 5(10), 813–819.

Fox, J., Friendly, G. G., Graves, S., Heiberger, R., Monette, G., Nilsson, H., … & Suggests, M. A. S. S. (2007). The car package. R Foundation for Statistical Computing.

Fuhrman, J. A. (2009). Microbial community structure and its functional implications. Nature, 459(7244), 193.

Galand, P. E., Casamayor, E. O., Kirchman, D. L., & Lovejoy, C. (2009). Ecology of the rare microbial biosphere of the Arctic Ocean. Proceedings of the National Academy of Sciences, 106(52), 22427–22432.

Gibbons, S. M., Caporaso, J. G., Pirrung, M., Field, D., Knight, R., & Gilbert, J. A. (2013). Evidence for a persistent microbial seed bank throughout the global ocean. Proceedings of the National Academy of Sciences, 110(12), 4651–4655.

Guerin, G. R., Sparrow, B., Tokmakoff, A., Smyth, A., Leitch, E., Baruch, Z., & Lowe, A. J. (2017). Opportunities for integrated ecological analysis across inland Australia with standardised data from Ausplots Rangelands. PloS one, 12(1), e0170137.

Hanson, C. A., Fuhrman, J. A., Horner-Devine, M. C., & Martiny, J. B. (2012). Beyond biogeographic patterns: processes shaping the microbial landscape. Nature Reviews Microbiology, 10(7), 497.

Hubbell, S. P. (2013). Tropical rain forest conservation and the twin challenges of diversity and rarity. Ecology and evolution, 3(10), 3263–3274.

Hubbell, S. P. (2001). The unified neutral theory of biodiversity and biogeography (MPB-32). Princeton University Press.

Hubbell, S. P. (1979). Tree dispersion, abundance, and diversity in a tropical dry forest. Science, 203(4387), 1299–1309.

Livermore, J. A., & Jones, S. E. (2015). Local–global overlap in diversity informs mechanisms of bacterial biogeography. The ISME journal, 9(11), 2413.

Locey, K. J., & Lennon, J. T. (2016). Scaling laws predict global microbial diversity. Proceedings of the National Academy of Sciences, 113(21), 5970–5975.

Logares, R., Sunagawa, S., Salazar, G., Cornejo‐Castillo, F. M., Ferrera, I., Sarmento, H., … & Raes, J. (2014). Metagenomic 16S rDNA I llumina tags are a powerful alternative to amplicon sequencing to explore diversity and structure of microbial communities. Environmental microbiology, 16(9), 2659–2671.

Louca, S., Mazel, F., Doebeli, M., & Parfrey, L. W. (2019). A census-based estimate of Earth’s bacterial and archaeal diversity. PLoS biology, 17(2), e3000106.

Lawless, J. F. (2003). Statistical Models and Methods for Life time Data, 3rd Ed. John Wiley and Sons, New York.

Loza, M. I., Jiménez, I., Jørgensen, P. M., Arellano, G., Macía, M. J., Torrez, V. W., & Ricklefs, R. E. (2017). Phylogenetic patterns of rarity in a regional species pool of tropical woody plants. Global ecology and biogeography, 26(9), 1043–1054.

Lynch, M. D., & Neufeld, J. D. (2015). Ecology and exploration of the rare biosphere. Nature Reviews Microbiology, 13(4), 217.

MacArthur, R. H. (1957). On the relative abundance of bird species. Proceedings of the National Academy of Sciences of the United States of America, 43(3), 293.

MacArthur, R. (1960). On the relative abundance of species. The American Naturalist, 94(874), 25–36.

Magurran, A. E., & Henderson, P. A. (2003). Explaining the excess of rare species in natural species abundance distributions. Nature, 422(6933), 714.

Martiny, J. B. H., Bohannan, B. J., Brown, J. H., Colwell, R. K., Fuhrman, J. A., Green, J. L., … & Morin, P. J. (2006). Microbial biogeography: putting microorganisms on the map. Nature Reviews Microbiology, 4(2), 102.

Matthews, T. J., Borregaard, M. K., Ugland, K. I., Borges, P. A., Rigal, F., Cardoso, P., & Whittaker, R. J. (2014). The gambin model provides a superior fit to species abundance distributions with a single free parameter: evidence, implementation and interpretation. Ecography, 37(10), 1002–1011.

Matthews, T. J., Borges, P. A., de Azevedo, E. B., & Whittaker, R. J. (2017). A biogeographical perspective on species abundance distributions: recent advances and opportunities for future research. Journal of biogeography, 44(8), 1705–1710.

Matthews, T. J., Sadler, J. P., Kubota, Y., Woodall, C. W., & Pugh, T. A. (2019). Systematic variation in North American tree species abundance distributions along macroecological climatic gradients. Global Ecology and Biogeography, 28(5), 601–611.

McGill, B. J., Etienne, R. S., Gray, J. S., Alonso, D., Anderson, M. J., Benecha, H. K., … & Hurlbert, A. H. (2007). Species abundance distributions: moving beyond single prediction theories to integration within an ecological framework. Ecology letters, 10(10), 995–1015.

McGill, B. J. (2003). Does Mother Nature really prefer rare species or are log‐left‐ skewed SADs a sampling artefact?. Ecology Letters, 6(8), 766–773.

Motomura, I. (1932). On the statistical treatment of communities. – Zoological Magazine (Tokyo), 44, 379–383.

Neves, D. M., Dexter, K. G., Pennington, R. T., Bueno, M. L., & Oliveira Filho, A. T. (2015). Environmental and historical controls of floristic composition across the South American Dry Diagonal. Journal of Biogeography, 42(8), 1566–1576.

Preston, F. W. (1948). The commonness, and rarity, of species. Ecology, 29(3), 254–283.

Quast, C., Pruesse, E., Yilmaz, P., Gerken, J., Schweer, T., Yarza, P., … & Glöckner, F. O. (2012). The SILVA ribosomal RNA gene database project: improved data processing and web-based tools. Nucleic acids research, 41(D1), D590–D596.

Quinn, G. P., & Keough, M. J. (2002). Experimental design and data analysis for biologists. Cambridge University Press.

Rabinowitz, D. (1981). Seven forms of rarity. Biological aspects of rare plant conservation.

Ricklefs, R. E. (2000). Rarity and diversity in Amazonian forest trees. Trends in Ecology & Evolution, 15(3), 83–84.

Rinne, H. (2008). The Weibull Distribution. A Handbook. CRC Press.

Rodriguez-R, L. M., Castro, J. C., Kyrpides, N. C., Cole, J. R., Tiedje, J. M., & Konstantinidis, K. T. (2018). How much do rRNA gene surveys underestimate extant bacterial diversity?. Appl. Environ. Microbiol., 84(6), e00014–18.

Rosindell, J., Hubbell, S. P., He, F., Harmon, L. J., & Etienne, R. S. (2012). The case for ecological neutral theory. Trends in ecology & evolution, 27(4), 203–208.

Schloss, P. D., Girard, R. A., Martin, T., Edwards, J., & Thrash, J. C. (2016). Status of the archaeal and bacterial census: an update. MBio, 7(3), e00201–16.

Ser-Giacomi, E., Zinger, L., Malviya, S., De Vargas, C., Karsenti, E., Bowler, C., & De Monte, S. (2018). Ubiquitous abundance distribution of non-dominant plankton across the global ocean. Nature ecology & evolution, 2(8), 1243.

Shade, A., Read, J. S., Youngblut, N. D., Fierer, N., Knight, R., Kratz, T. K., … & Whitaker, R. J. (2012). Lake microbial communities are resilient after a whole-ecosystem disturbance. The ISME journal, 6(12), 2153.

Shade, A., Dunn, R. R., Blowes, S. A., Keil, P., Bohannan, B. J., Herrmann, M., … & Chase, J. (2018). Macroecology to unite all life, large and small. Trends in ecology & evolution.

Sherrill-Mix, S., McCormick, K., Lauder, A., Bailey, A., Zimmerman, L., Li, Y., … & Hart, T. B. (2018). Allometry and ecology of the bilaterian gut microbiome. MBio, 9(2), e00319–18.

Shoemaker, W. R., Locey, K. J., & Lennon, J. T. (2017). A macroecological theory of microbial biodiversity. Nature ecology & evolution, 1(5), 0107.

Sogin, M. L., Morrison, H. G., Huber, J. A., Welch, D. M., Huse, S. M., Neal, P. R., … & Herndl, G. J. (2006). Microbial diversity in the deep sea and the underexplored “rare biosphere”. Proceedings of the National Academy of Sciences, 103(32), 12115–12120.

Stauffer, H. B. (1979). A derivation for the Weibull distribution. Journal of theoretical biology, 81(1), 55–63.

Stegen, J. C., Hurlbert, A. H., Bond-Lamberty, B., Chen, X., Anderson, C. G., Chu, R. K., … & Tfaily, M. (2016). Aligning the measurement of microbial diversity with macroecological theory. Frontiers in microbiology, 7, 1487.

Sunagawa, S., Coelho, L. P., Chaffron, S., Kultima, J. R., Labadie, K., Salazar, G., … & Cornejo-Castillo, F. M. (2015). Structure and function of the global ocean microbiome. Science, 348(6237), 1261359.

Tabachnick, B. G., Fidell, L. S., & Ullman, J. B. (2007). Using multivariate statistics (Vol. 5). Boston, MA: Pearson.

Ugland, K. I., Lambshead, P. J. D., McGill, B., Gray, J. S., O’Dea, N., Ladle, R. J., & Whittaker, R. J. (2007). Modelling dimensionality in species abundance distributions: description and evaluation of the Gambin model. Evolutionary Ecology Research, 9(2), 313–324.

Ulrich, W., & Ollik, M. (2004). Frequent and occasional species and the shape of relative‐abundance distributions. Diversity and distributions, 10(4), 263–269.

Ulrich, W., Kusumoto, B., Shiono, T., & Kubota, Y. (2016a). Climatic and geographic correlates of global forest tree species–abundance distributions and community evenness. Journal of vegetation science, 27(2), 295–305.

Ulrich, W., Soliveres, S., Thomas, A. D., Dougill, A. J., & Maestre, F. T. (2016b). Environmental correlates of species rank−abundance distributions in global drylands. Perspectives in plant ecology, evolution and systematics, 20, 56–64.

Ulrich, W., Nakadai, R., Matthews, T., Kubota, Y., (2018a). https://figshare.com/articles/Weibull_fits/5975098.

Ulrich, W., Nakadai, R., Matthews, T. J., & Kubota, Y. (2018b). The two-parameter Weibull distribution as a universal tool to model the variation in species relative abundances. Ecological complexity, 36, 110–116.

Ulrich, W., Ollik, M., & Ugland, K. I. (2010). A meta‐analysis of species–abundance distributions. Oikos, 119(7), 1149–1155.

Vellend, M. (2010). Conceptual synthesis in community ecology. The Quarterly review of biology, 85(2), 183–206.

Vellend, M. (2016). The theory of ecological communities (MPB-57) (Vol. 75). Princeton University Press.

White, E. P., Thibault, K. M., & Xiao, X. (2012). Characterizing species abundance distributions across taxa and ecosystems using a simple maximum entropy model. Ecology, 93(8), 1772–1778.

Weibull, W., 1951. A statistical distribution function of wide applicability. Journal of Applied Mechanics 18, 239–296.

Whittaker, R.H. (1975). Communities and Ecosystems, 2nd edn. MacMillan, New York NY, US.

Zhang, D. (2018). rsq: R-squared and related measures. R package version, 1(1).

