## Supplementary figures and images for "Describing macroecological patterns in microbes: Approaches for comparative analyses of operational taxonomic unit read number distribution with a case study of global oceanic bacteria"

### Figure S1

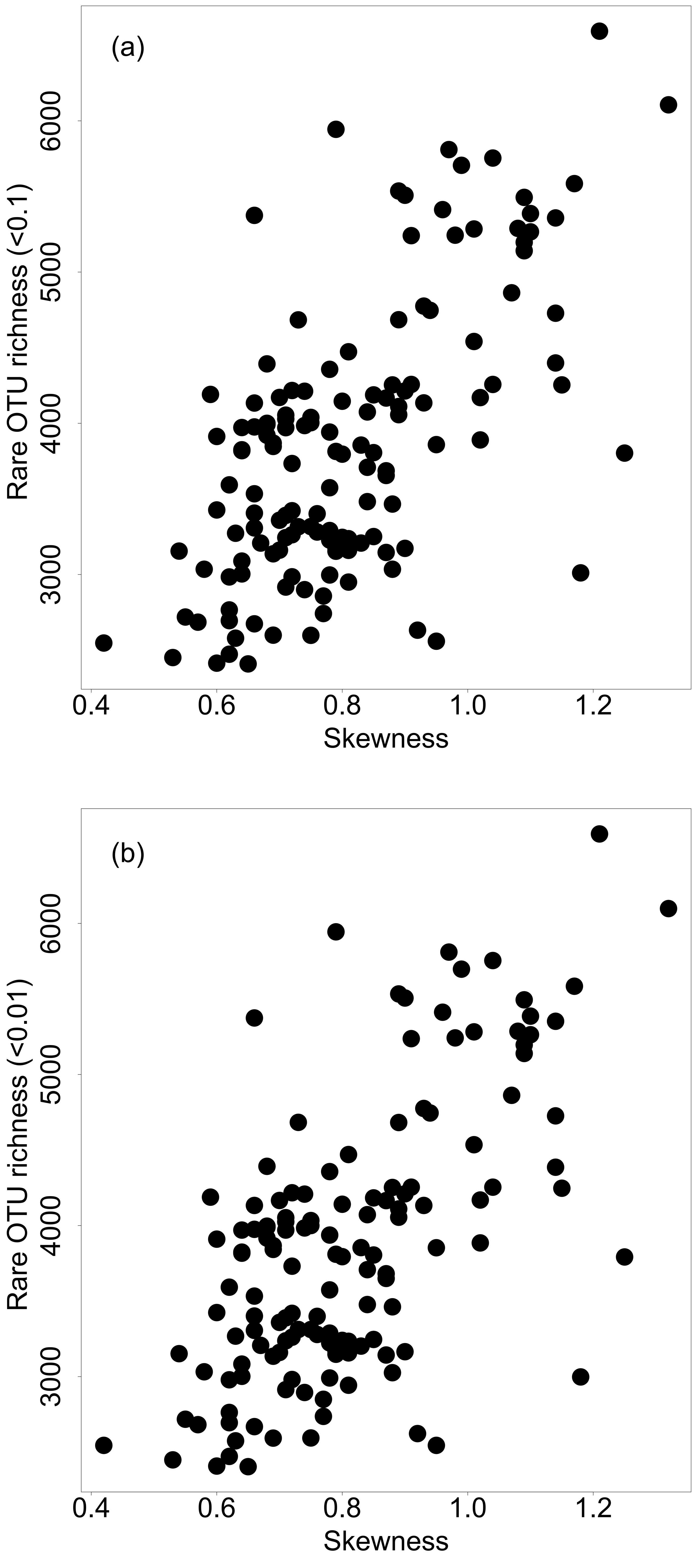

### Figure S2

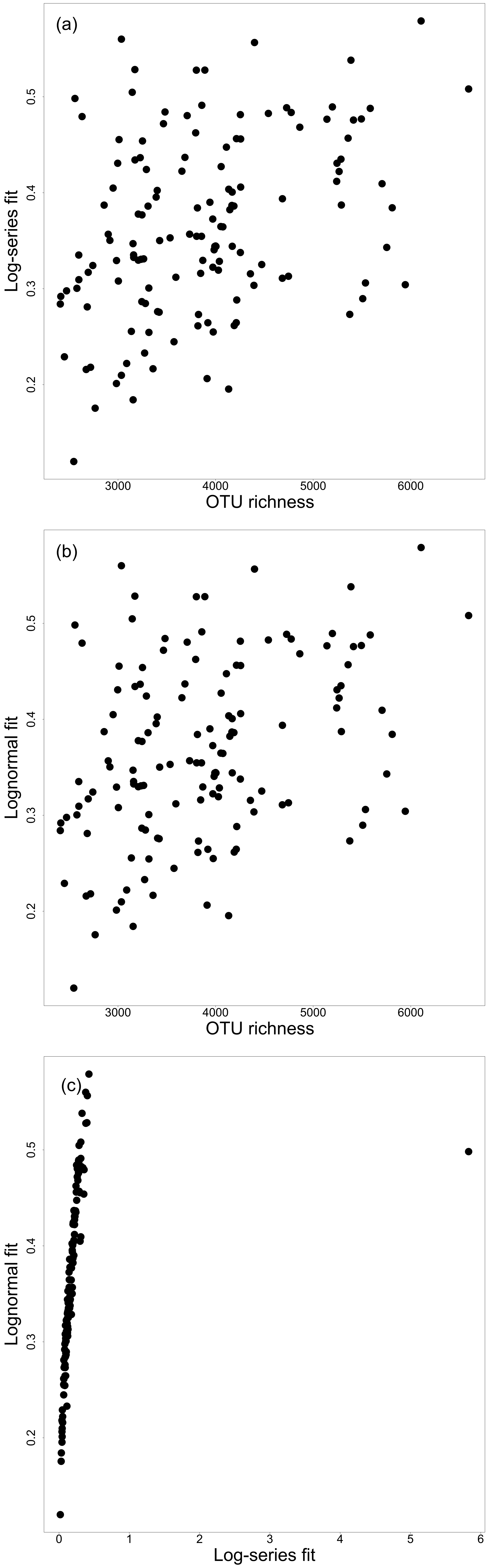

### Figure S3

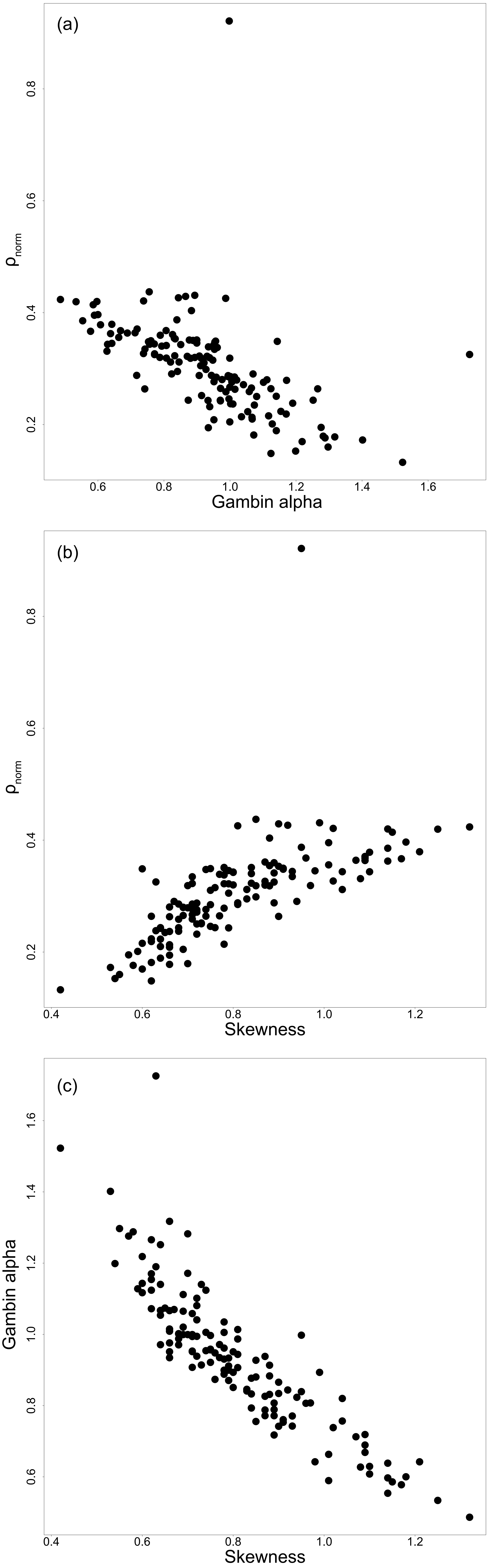

### Figure S4

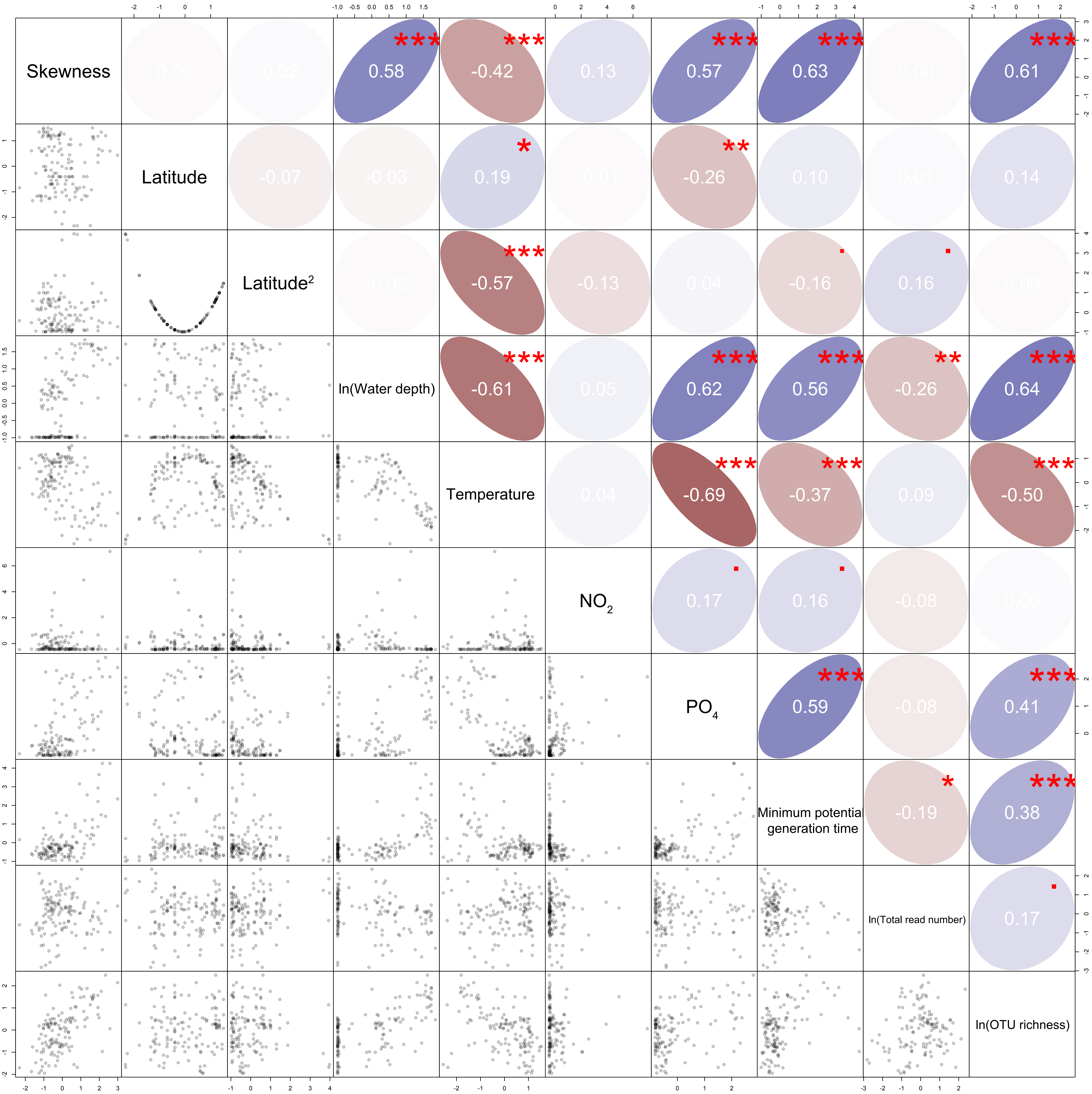
