## Supplemental text 1 for "Describing macroecological patterns in microbes: Approaches for comparative analyses of operational taxonomic unit read number distribution with a case study of global oceanic bacteria"

**Supplementary text 1 Multimodality of OTU read number distributions**

Datasets containing multi-property groups have been discussed in the context of multimodality (Mathews *et al*. 2019 a, b). As a preliminary analysis, we fitted microbial ORDs to a multi-modal gambin distribution and compared the modal numbers of the best fitting models using the Bayesian information criterion (BIC). The results clearly supported the three-modal distribution as the best fit for ORDs of bacterial communities (123/139: 88.5%), and no model supported the unimodal distribution as the best fit. Mathews *et al*. (2019a) reported that only 19% (51 of 275 datasets) of tree communities follow a bimodal distribution. Therefore, bacterial communities have a greater tendency for multimodality than tree communities. These differences in multimodality may be related to the properties of high-throughput sequencing data containing multi-trophic OTUs. Here, we were unable to explore the detailed mechanisms of multimodality in ORDs because multimodality and its ecological mechanisms remain poorly understood, even in macroscopic organisms (Mathews *et al*. 2019a). Further studies are needed, such as simulation studies focusing on the microbial assemblage to identify the parameters with ecological and mechanistic meaning, similar to Hubbell (2001).

|  | Mode number | | | | |
| --- | --- | --- | --- | --- | --- |
|  | 1 | 2 | 3 | 4 | 5 |
| b_1 | 14968.3348 | 14845.3972 | **14768.7915** | 14803.4766 | 14816.6078 |
| b_2 | 15662.3248 | 15527.7733 | **15455.4542** | 15504.3530 | 15502.5925 |
| b_3 | 11647.2136 | 11567.3795 | **11503.5448** | 11529.9171 | 11533.8496 |
| b_4 | 12693.0673 | 12609.4056 | **12569.3399** | 12584.1353 | 12595.7483 |
| b_5 | 15378.0708 | 15249.1087 | **15129.6787** | 15145.4760 | 15143.5008 |
| b_6 | 15670.4881 | 15548.6479 | **15524.8898** | 15558.4662 | 15555.5209 |
| b_7 | 13949.9299 | 13861.3973 | **13847.0524** | 13880.4566 | 13920.462 |
| b_8 | 11728.5080 | 11614.2608 | **11548.2854** | 11571.0221 | 11573.1536 |
| b_9 | 12748.5861 | 12615.3357 | **12584.4945** | 12588.3356 | 12615.9336 |
| b_10 | 16507.7533 | 16422.5722 | **16388.4956** | 16410.8763 | 16419.6189 |
| b_11 | 10458.3360 | 10364.5219 | **10338.7925** | 10362.5501 | 10366.6737 |
| b_12 | 9691.05824 | 9579.46588 | **9520.6231** | 9570.25647 | 9603.39849 |
| b_13 | 12651.4084 | 12581.2433 | **12558.8828** | 12569.1328 | 12580.1769 |
| b_14 | 9759.4209 | 9612.3223 | **9598.6062** | 9634.2491 | 9647.1593 |
| b_15 | 12775.3516 | 12685.7823 | **12637.5764** | 12656.3093 | 12669.2074 |
| b_16 | 9542.9486 | 9452.7874 | **9408.3107** | 9440.6923 | 9447.9056 |
| b_17 | 15027.3231 | 14827.0853 | **14790.6592** | 14796.8027 | 14808.3869 |
| b_18 | 11990.5861 | 11885.2060 | **11881.5332** | 11882.9589 | 11901.9575 |
| b_19 | 10377.5849 | 10271.7813 | **10258.4476** | 10260.8860 | 10269.6517 |
| b_20 | 9898.2712 | 9783.1477 | **9776.3164** | 9783.4267 | 9810.0228 |
| b_21 | 10263.0119 | 10097.8673 | **10094.8762** | 10116.4014 | 10123.4331 |
| b_22 | 13022.8675 | 12833.8840 | **12817.4299** | 12827.2329 | 12833.0215 |
| b_23 | 11940.7083 | **11858.0385** | 11858.8573 | 11876.4050 | 11880.9177 |
| b_24 | 13291.5680 | **13143.3977** | 13148.5775 | 13158.7568 | 13167.7055 |
| b_25 | 10258.0822 | 10156.3659 | **10137.7409** | 10152.1154 | 10162.4445 |
| b_26 | 12156.7843 | **11977.5421** | 11985.1542 | 12002.6651 | 12008.054 |
| b_27 | 12481.9752 | **12305.4690** | 12312.6664 | 12319.3785 | 12334.4006 |
| b_28 | 12152.3893 | 11931.1181 | **11928.4249** | 11966.5142 | 11999.1253 |
| b_29 | 17181.1146 | 17023.2983 | **16927.0717** | 16958.3333 | 16945.5569 |
| b_30 | 13560.2472 | 13456.7554 | **13353.5882** | 13371.7666 | 13394.0476 |
| b_31 | 12941.4359 | 12825.7095 | **12793.8825** | 12821.6564 | 12839.5738 |
| b_32 | 11490.8015 | 11396.0813 | **11359.2498** | 11390.2125 | 11403.3728 |
| b_33 | 14377.9053 | 14247.5656 | 14209.7933 | **14208.3373** | 14218.8189 |
| b_34 | 12095.8799 | 11987.4510 | **11952.7505** | 11972.1404 | 11973.2070 |
| b_35 | 15494.4490 | 15383.4213 | **15293.4163** | 15310.7244 | 15328.4146 |
| b_36 | 16606.6742 | 16497.1776 | **16442.6780** | 16458.8343 | 16477.3237 |
| b_37 | 15736.9977 | 15588.7164 | **15522.7556** | 15570.6476 | 15569.7811 |
| b_38 | 15905.9448 | 15792.4998 | **15762.2500** | 15773.0437 | 15802.2764 |
| b_39 | 13737.5003 | 13611.3419 | **13545.8856** | 13622.9309 | 13634.0180 |
| b_40 | 15666.9701 | 15532.0729 | **15489.9773** | 15527.8290 | 15507.4083 |
| b_41 | 11717.2366 | 11607.3530 | **11523.3822** | 11542.5537 | 11553.2774 |
| b_42 | 10811.8142 | 10641.2823 | **10624.1508** | 10668.8394 | 10675.5582 |
| b_43 | 11892.3280 | 11763.5944 | **11685.8790** | 11704.4471 | 11757.5654 |
| b_44 | 16996.1019 | 16892.5969 | **16835.3187** | 16878.3934 | 16850.4037 |
| b_45 | 13591.2968 | 13396.2515 | **13303.3808** | 13393.0956 | 13405.7015 |
| b_46 | 16073.9560 | 15872.8956 | **15810.1302** | 15888.2289 | 15881.9813 |
| b_47 | 20413.3035 | 20196.7347 | **20016.4791** | 20085.1144 | 20094.4134 |
| b_48 | 14116.4756 | 13902.1149 | **13878.7597** | 13902.6045 | 13909.4788 |
| b_49 | 14756.4700 | 14584.9564 | **14535.2505** | 14601.3274 | 14602.9740 |
| b_50 | 15778.2495 | 15619.7086 | **15560.9318** | 15584.9401 | 15628.3071 |
| b_51 | 13628.1408 | 13455.0823 | **13368.9406** | 13399.3055 | 13446.6015 |
| b_52 | 15628.6613 | 15461.8147 | **15442.5656** | 15459.4093 | 15464.4788 |
| b_53 | 17879.2911 | 17710.1448 | **17643.4829** | 17659.0975 | 17670.7076 |
| b_54 | 14744.4559 | 14629.1447 | **14627.7615** | 14645.3714 | 14656.0170 |
| b_55 | 16099.7368 | 15956.7219 | **15936.0888** | 15949.1155 | 15971.2425 |
| b_56 | 21715.3672 | 21555.6663 | **21484.7045** | 21526.4208 | 21526.2523 |
| b_57 | 13000.1752 | 12818.6433 | **12815.7101** | 12821.0790 | 12845.9111 |
| b_58 | 19031.6244 | 18855.3389 | 18807.0275 | **18805.2916** | 18814.3716 |
| b_59 | 13657.2275 | 13514.5751 | **13453.9265** | 13473.0408 | 13513.1945 |
| b_60 | 19639.1372 | 19461.2762 | **19386.9032** | 19402.4337 | 19434.4058 |
| b_61 | 15154.8258 | 14960.5875 | **14916.5628** | 14935.1366 | 14956.9731 |
| b_62 | 14825.4211 | 14747.7104 | **14732.8410** | 14753.0672 | 14756.7497 |
| b_63 | 14562.4566 | 14342.9746 | **14310.5433** | 14349.6301 | 14392.0780 |
| b_64 | 14083.6603 | 13978.3631 | **13970.2944** | 13988.3045 | 13992.1076 |
| b_65 | 15228.6309 | 15098.1576 | **15042.2426** | 15076.8303 | 15090.3829 |
| b_66 | 17114.5229 | 16976.5206 | 16916.7941 | **16912.7395** | 16917.6600 |
| b_67 | 18725.0128 | 18557.8270 | **18510.4450** | 18535.4976 | 18551.4560 |
| b_68 | 11959.4791 | 11792.2860 | **11772.0938** | 11798.3050 | 11830.4152 |
| b_69 | 16394.1145 | 16262.1905 | **16246.9125** | 16263.2805 | 16268.5193 |
| b_70 | 22100.0266 | 21980.2361 | **21968.6696** | 21995.7182 | 21983.9177 |
| b_71 | 16916.5621 | 16771.7658 | **16709.5311** | 16732.1673 | 16759.3800 |
| b_72 | 15883.9395 | 15756.9328 | **15723.3586** | 15734.5466 | 15745.7468 |
| b_73 | 20531.7990 | 20277.8642 | 20253.8444 | **20252.2141** | 20262.7080 |
| b_74 | 13103.1452 | 12875.4567 | **12837.5881** | 12899.2259 | 12901.3987 |
| b_75 | 12912.1491 | 12796.3733 | **12718.2097** | 12733.2346 | 12742.0500 |
| b_76 | 18570.1228 | 18288.8346 | **18251.3469** | 18253.1239 | 18255.4480 |
| b_77 | 14491.0609 | 14359.0600 | **14354.5058** | 14375.8293 | 14397.1095 |
| b_78 | 16063.0985 | 15918.5469 | **15890.0996** | 15897.9573 | 15948.8587 |
| b_79 | 13601.4442 | 13467.4401 | **13416.8773** | 13422.6108 | 13454.7128 |
| b_80 | 15644.8102 | 15494.9593 | **15493.1907** | 15527.4256 | 15545.8325 |
| b_81 | 18763.7670 | 18573.0057 | **18478.5706** | 18484.4877 | 18518.9765 |
| b_82 | 15501.0719 | 15324.6517 | 15280.9434 | **15273.9061** | 15327.3420 |
| b_83 | 18744.8909 | 18606.0626 | **18563.0781** | 18583.1202 | 18581.4817 |
| b_84 | 14943.8514 | 14847.5908 | **14766.4028** | 14784.2509 | 14789.3186 |
| b_85 | 15905.9755 | 15792.4826 | **15756.3859** | 15793.3141 | 15802.6370 |
| b_86 | 15553.4499 | 15407.3855 | **15350.6767** | 15400.5126 | 15411.2781 |
| b_87 | 17889.9241 | 17746.6157 | **17686.6953** | 17706.8038 | 17720.9740 |
| b_88 | 15608.8908 | 15392.1026 | **15315.8960** | 15346.9021 | 15390.9146 |
| b_89 | 17985.7029 | 17850.5524 | **17833.7209** | 17879.8576 | 17880.2909 |
| b_90 | 15735.4987 | 15522.3673 | **15417.8069** | 15501.9633 | 15512.5284 |
| b_91 | 15809.9234 | 15611.0881 | **15534.4614** | 15621.9995 | 15618.2421 |
| b_92 | 15960.7179 | 15796.1683 | **15658.8106** | 15713.2711 | 15708.5224 |
| b_93 | 17741.0037 | 17555.1304 | **17473.1734** | 17484.5670 | 17485.2617 |
| b_94 | 15451.7767 | 15296.0089 | **15247.3133** | 15310.0066 | 15310.2682 |
| b_95 | 17120.4942 | 16952.6637 | 16875.2166 | **16881.6100** | 16906.2872 |
| b_96 | 18040.7391 | 17923.2629 | **17884.2518** | 17905.8625 | 17931.7594 |
| b_97 | 14914.7493 | 14798.4411 | **14699.7079** | 14735.6363 | 14726.6053 |
| b_98 | 19873.4699 | 19504.1083 | 19476.7189 | **19468.5438** | 19505.2202 |
| b_99 | 12673.2822 | 12458.1330 | **12439.4903** | 12469.1976 | 12473.3157 |
| b_100 | 13330.7772 | 13105.9933 | **13084.8965** | 13091.3989 | 13099.4043 |
| b_101 | 12608.9447 | 12420.2651 | **12414.4828** | 12463.5764 | 12450.4538 |
| b_102 | 10678.5853 | 10488.7733 | **10481.6801** | 10520.4063 | 10525.2056 |
| b_103 | 15014.1188 | 14829.7885 | **14819.0866** | 14838.9681 | 14848.6162 |
| b_104 | 13995.1369 | 13858.5265 | **13845.6591** | 13860.2754 | 13874.1181 |
| b_105 | 14600.3200 | 14384.9590 | **14336.4677** | 14398.6639 | 14393.8665 |
| b_106 | 20347.8408 | 20203.6360 | 20198.6853 | **20192.6687** | 20219.9504 |
| b_107 | 19153.7860 | 19027.5839 | **18997.4675** | 19020.9365 | 19072.2155 |
| b_108 | 10644.7605 | 10550.8949 | **10491.0081** | 10543.4586 | 10553.4614 |
| b_109 | 10138.6937 | 10003.2922 | **10000.2723** | 10008.4963 | 10010.4865 |
| b_110 | 12370.0165 | 12193.2856 | **12174.7149** | 12183.1152 | 12205.0840 |
| b_111 | 11888.8240 | **11813.1379** | 11814.9464 | 11819.1716 | 11832.1629 |
| b_112 | 11387.7779 | 11246.6172 | **11209.1662** | 11220.4393 | 11231.8915 |
| b_113 | 17332.2504 | 17073.8452 | **17069.1404** | 17076.3261 | 17132.5794 |
| b_114 | 12563.0620 | 12445.0432 | **12438.3090** | 12444.8666 | 12458.1113 |
| b_115 | 16615.7201 | 16372.5451 | 16271.5460 | **16264.4695** | 16280.7169 |
| b_116 | 9866.70798 | 9661.95354 | **9637.95891** | 9652.39634 | 9665.07096 |
| b_117 | 11575.2519 | 11486.2320 | **11420.5986** | 11438.2387 | 11453.7149 |
| b_118 | 19217.3501 | 18974.6274 | **18888.6509** | 18896.8890 | 18912.1474 |
| b_119 | 12838.8196 | 12736.1329 | **12724.8185** | 12744.4686 | 12769.0611 |
| b_120 | 16597.0414 | 16501.6810 | **16475.8258** | 16484.8820 | 16486.7329 |
| b_121 | 20600.6625 | 20477.0215 | **20431.5767** | 20470.9955 | 20489.9667 |
| b_122 | 10578.0408 | 10450.9562 | **10402.0889** | 10428.4619 | 10436.3068 |
| b_123 | 15859.6577 | 15782.5268 | **15761.7815** | 15784.6581 | 15799.5029 |
| b_124 | 19359.7470 | **19272.4402** | 19273.3913 | 19284.3368 | 19301.4077 |
| b_125 | 12168.2993 | 12096.4215 | **12040.3509** | 12068.5310 | 12082.7996 |
| b_126 | 16449.7982 | 16303.1433 | **16251.2240** | 16269.2945 | 16277.1620 |
| b_127 | 12191.4505 | 12024.7292 | **12017.4302** | 12028.5243 | 12032.3282 |
| b_128 | 12032.7204 | 11955.6768 | **11945.7460** | 11965.3954 | 11974.0578 |
| b_129 | 10612.7435 | 10555.3440 | **10541.7484** | 10557.5871 | 10563.6779 |
| b_130 | 16924.8841 | 16815.1343 | **16790.8736** | 16813.2155 | 16821.5252 |
| b_131 | 10548.5137 | **10515.4304** | 10522.9544 | 10530.3075 | 10541.6138 |
| b_132 | 22163.9143 | **22074.1016** | 22074.3644 | 22081.2695 | 22090.7191 |
| b_133 | 16026.9088 | 15937.6496 | **15904.6279** | 15911.6704 | 15921.6451 |
| b_134 | 15569.0210 | 15450.4193 | **15370.1157** | 15394.8362 | 15412.0777 |
| b_135 | 13065.3510 | 12949.6451 | **12858.6320** | 12881.6528 | 12905.0973 |
| b_136 | 12245.5308 | 12161.1297 | **12124.7981** | 12137.6041 | 12145.9333 |
| b_137 | 15015.3566 | 14890.8389 | **14841.5341** | 14848.2618 | 14855.1619 |
| b_138 | 10231.5021 | 10151.9731 | **10083.8234** | 10104.2099 | 10126.6024 |
| b_139 | 11626.0063 | 11528.3108 | **11467.7064** | 11515.8225 | 11517.0792 |

*The values indicate BIC values.

**References**

Hubbell, S. P. (2001). The unified neutral theory of biodiversity and biogeography (MPB-32). Princeton University Press.

Matthews, T. J., Borregaard, M. K., Gillespie, C. S., Rigal, F., Ugland, K. I., Krüger, R. F., ... & Whittaker, R. J. (2019a). Extension of the gambin model to multimodal species abundance distributions. Methods in Ecology and Evolution, 10(3), 432-437.

Matthews, T. J., Sadler, J. P.,Kubota, Y., Woodall, C. W., & Pugh, T. A. (2019b). Systematic variation in North American tree species abundance distributions along macroecological climatic gradients. Global Ecology and Biogeography, 28(5), 601-611.
